# Disulfide bond sculpts a peptide fold that mediates phytocytokine recognition

**DOI:** 10.1101/2025.10.24.684422

**Authors:** Pedro Jiménez-Sandoval, Oliver Johanndrees, Simon Snoeck, Chitthavalli Y. Harshith, Moutassem Omary, Caroline Broyart, Jack Rhodes, Kyle W. Bender, Cyril Zipfel, Julia Santiago

## Abstract

Precise ligand recognition by closely related leucine-rich repeat receptor kinases (LRR-RKs) is essential for plants to coordinate immunity, development, and environmental adaptation. Here, we show how HAESA-LIKE 3 (HSL3) specifically recognizes the folded, disulfide-stabilized CTNIP/SCREW phytocytokines in Arabidopsis. Quantitative binding assays define a minimal CTNIP4 region required for high-affinity HSL3 interaction and signaling activation. A 2.12 Å crystal structure of the HSL3-CTNIP4 complex reveals a unique C-terminal receptor pocket that accommodates the peptide’s cyclic architecture through a combination of hydrophobic and polar contacts, a feature absent in the closely related HAE/HSL LRR-RKs. The cyclic CTNIP4 fold further establishes a largely hydrophobic interface that bridges HSL3 to the SERK co-receptor, forming a distinct activation surface. Together, these structural, biochemical and physiological insights uncover a previously unrecognised mechanism of peptide perception and receptor activation, highlighting how subtle architectural variations enable precise ligand selectivity among highly conserved plant receptor kinases.

## Main

Plant peptide hormones have emerged as central mediators of cell-to-cell communication, coordinating growth, development, and environmental adaptation such as immune responses^1–14^. These signaling molecules are typically processed from larger precursor proteins and undergo diverse post-translational modifications including tyrosine sulfation, proline hydroxylation, glycosylation or the formation of disulfide bonds^3,6,7,12,15–20^. Such chemical modifications can critically influence peptide diffusion, stability, and bioactivity, enabling precise recognition by their cognate cell-surface receptors ^6,7,9^. To perceive an extensive repertoire of sequence-diverse signaling peptides, plants have co-evolved an equally diverse array of cell-surface localized receptors. Among these, leucine-rich repeat receptor kinases (LRR-RKs) represent the largest and most functionally versatile family^21^. Despite their structural similarity, individual LRR-RKs have diversified their extracellular binding pockets to prevent ligand cross-reactivity and ensure specific recognition of their *bona fide* peptide hormones^6^. Typically, LRR-RKs initiate a specific signalling response through the ligand-dependent recruitment of a shape-complementary co-receptor, forming an active receptor complex at the plasma membrane^7–9,21–23^.

Recently, we and others identified a new family of plant signaling peptides acting as phytocytokines, termed CTNIPs or SMALL PHYTOCYTOKINES REGULATING DEFENSE AND WATER LOSS (SCREWs), together with their cognate receptor, the leucine-rich repeat receptor kinase HAESA-LIKE 3 (HSL3), also known as PLANT SCREW UNRESPONSIVE RECEPTOR (NUT)^1,4,24^. CTNIP/SCREW peptides contain a pair of conserved cysteine residues that are essential for their bioactivity and for recognition by HSL3/NUT^4^. Ligand binding promotes heteromerization of HSL3/NUT with members of the SOMATIC EMBRYOGENESIS RECEPTOR-LIKE KINASE (SERK) co-receptor family such as SERK3 (also named BRASSINOSTEROID INSENSITIVE 1-ASSOCIATED KINASE 1, BAK1), forming an active receptor complex that triggers stress-induced responses^1,4^. However, despite its distal homology to the HAESA (HAE) and HAESA-LIKE 1/2 (HSL1/2) receptors, HSL3 features a distinct ligand-binding pocket that lacks the conserved motifs required for INFLORESCENCE DEFICIENT IN ABSCISSION (IDA)/IDA-LIKE (IDL) peptide recognition^4,6,7^, suggesting that it employs a novel mechanism to perceive CTNIP/SCREW peptides. Elucidating this unique mode of peptide recognition and receptor activation is the central focus of this study.

Building on our previous identification of CTNIP peptides and the report of micromolar range binding affinities^4^, we next sought to define a minimal CTNIP region required for high-affinity (nanomolar range) recognition by the HSL3 receptor. To this end, we synthetized multiple longer or shorter N-terminal versions of CTNIP4 (corresponding to SCREW2) and assessed binding of the resulting peptide variants to the HSL3 ectodomain using isothermal titration calorimetry (ITC) (Fig. 1a and Supplementary Fig. 1). Given the essential role of the conserved cysteine residues for receptor interaction^4^, all variants were synthesised with a disulfide bond linking Cys58 and Cys68 to stabilise the folded conformation and peptide homogeneity (see Methods). ITC analyses revealed that CTNIP4(46-70)_Cys-Cys_ exhibited the highest binding affinity to HSL3, defining the peptide length most effectively coordinated by the receptor-binding pocket. The interaction is enthalpy-driven, indicating that peptide-receptor recognition is dominated by specific polar and hydrogen-bonding interactions rather than by hydrophobic effects (Supplementary Fig. 2). In contrast, longer CTNIP4 versions displayed up to a 60-fold reduction in affinity, suggesting that additional N-terminal residues may hinder receptor recognition. Stepwise truncation further refined this interaction: removal of one residue CTNIP4(47-70)_Cys-Cys_ resulted in a ∼4-fold reduction in binding, while deletion of two residues, CTNIP4(48-70)_Cys-Cys_, led to a dramatic ∼200-fold loss of affinity (Fig. 1b and Supplementary Fig. 1 and 2). These results identify Gln46 and Arg47 as key anchoring residues required for optimal receptor-peptide coordination. Consistent with these observations, the linear (non-cyclised) CTNIP4^C58A.C68A^(46-70) and the shorter and non-folded CTNIP4(48-70)_linear_ peptides, displayed either severely impaired (∼200-fold) or non detectable binding, respectively. These results highlight the critical contribution of both the N-ter anchoring residues and the disulfide-stabilised cyclic conformation to high affinity and thermodynamically favourable receptor-peptide association (Fig. 1b and Supplememtary Fig. 2). To assess whether binding affinities correlate with biological activity, we next tested the CTNIP4 variants for their ability to trigger downstream signaling responses in planta. Measurements of cytosolic Ca^2+^ influx and MAP kinase activation closely mirrored the in vitro binding profiles, with only peptides retaining high-affinity interaction with HSL3 inducing robust signalling responses (Fig. 1c,d and Supplementary Fig. 2). Together, these results delineate the core structural features and sequence determinants of CTNIP4 required for receptor recognition and signaling activation.

**Fig. 1:**
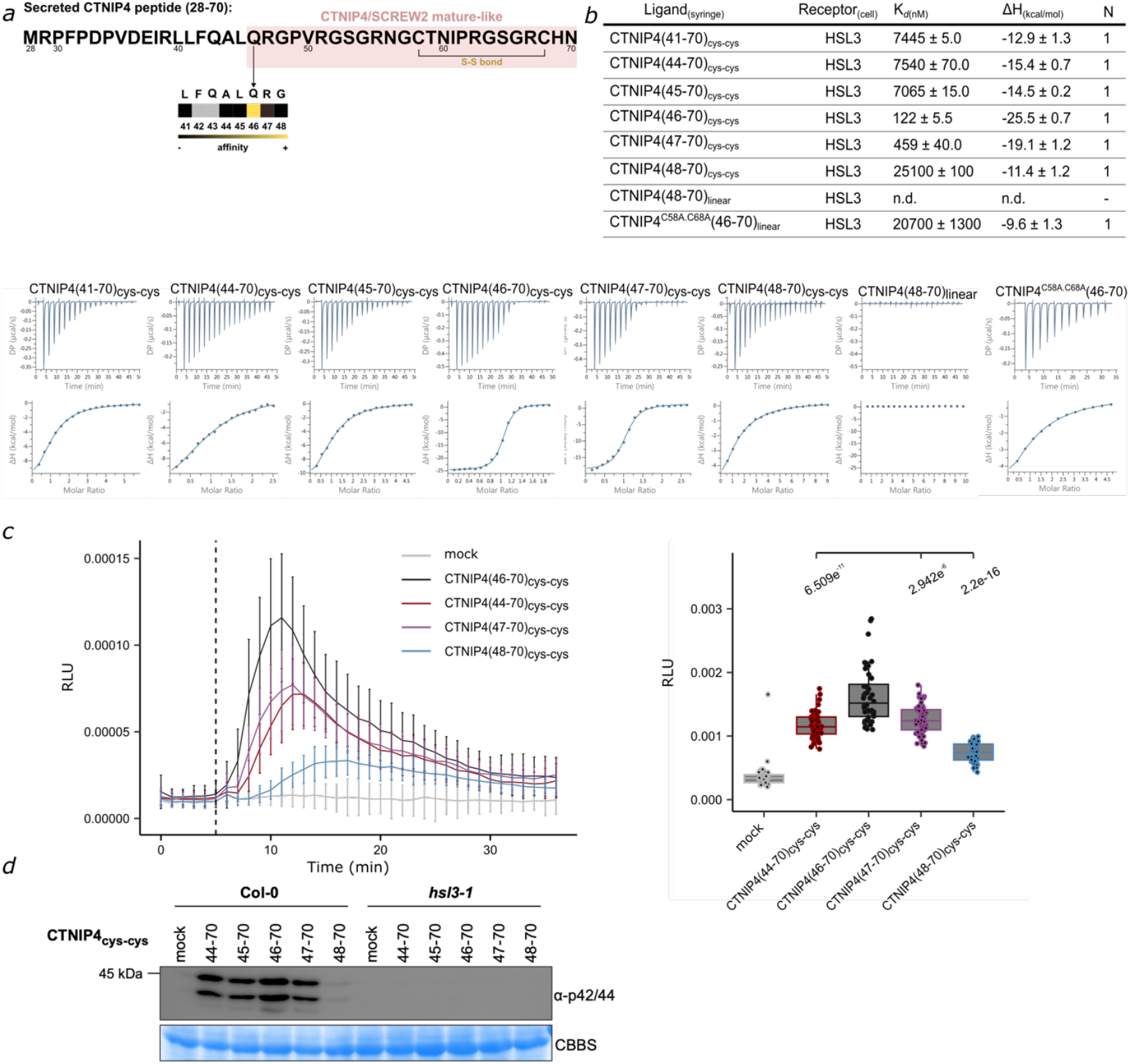
Mapping of a bioactive CTNIP4/SCREW2 peptide specifically recognized by the HSL3 receptor. Systematic biochemical analysis defines the minimal region of the CTNIP4/SCREW2 peptide required for high-affinity recognition by the HSL3 receptor. **a**, Schematic representation of the CTNIP4 secreted precursor. The N-terminal truncations used for biochemical analysis are indicated below. The minimal region required for high-affinity binding, including the conserved disulfide bond, is highlighted in pink. **b**, Isothermal titration calorimetry (ITC) assays showing binding of HSL3 ectodomain to folded and linear CTNIP4 peptides of varying lengths. ITC table summarizes Kd (nM), stoichiometry (N=1); n.d., no binding detected. Data are presented as mean ± SD from at least two independent experiments. **c**, Ca^+2^ influx assays assays in relative luminescence units (RLUs) show that folded CTNIP4(46-70)_cys-cys_ elicit the highest response in wild-type Col-0^AEQ^ plants compared to other lengths. Peptide concentration used was100 nM. Data represents mean ± SD (n=48), the vertical dotted line indicates the timing of the treatment. Considering the cumulative RLUs (3-30 min post treatment), the box plots indicate the median and the interquartile range (IQR), the whiskers extend to the most extreme data points within 1.5 times the IQR. Significance was tested by performing non-parametric Wilcoxon-Mann-Whitney tests between CTNIP4(46-70)_cys-cys_ and CTNIP4 length variants. **d**, Immunoblot analysis of MAP kinase activation in response to folded 5 nM CTNIP4 peptides of different lengths for 20 min, confirming that functional activity correlates with high-affinity receptor binding. Data shown are from one representative experiment out of three that yielded similar results.

To elucidate the molecular basis of CTNIP4 recognition, we determined the crystal structure of the HSL3 ectodomain in complex with the bioactive CTNIP4(46-70)_Cys-Cys_ peptide at 2.12 Å resolution (Supplementary Table 1). The complex crystalized in a 1:1 stoichoimetry arrangement (Fig. 2a and Supplementary Fig. 3), with the receptor extracellular adopting the canonical architecture of that of a LRR-RK. The HSL3 ectodomain comprises 21 LRRs flanked by N- and C-terminal capping domains that frame a slightly curved solenoid scaffold (Supplementary Fig. 4)^25^. The structure reveals that CTNIP4 adopts a unique disulfide-stabilised cyclic fold that is deeply embedded within the C-terminal portion of the HSL3 ligand-binding groove (Fig. 2a and Supplementary Fig. 3). This compact, looped conformation is sculpted by a peptide intramolecular hydrogen-bond network and the Cys58-Cys68 disulfide bond, which locks the peptide into a three-dimensional architecture optimised for receptor recognition (Supplementary Fig. 3). B-factor analysis indicates a gradient of coordination along the peptide: the linear N-terminal and core region of the peptide, extending up to Arg63, are highly ordered and tightly coordinated within the receptor pocket, whereas the cyclic loop region shows higher flexibility until the segment encompassing the disulfide bond and His69, where the covalent linkage restricts peptide flexibility and stabilises the fold (Fig. 2b). The peptide preorganized conformation facilitates its precise accommodation within the receptor cavity. Consistent with this structural model, the truncated linear variant CTNIP4^C58A^(46–63), which includes only the highly coordinated region of the peptide but lacks both the C-terminal loop and the disulfide bond constraint, exhibited an ∼ 200-fold reduction in binding affinity to HSL3 (Fig. 2c). These data reinforce the functional importance of the C-terminal cyclic loop, whose conformational rigidity and defined topology are critical for high-affinity and specific receptor recognition. Detailed inspection of the receptor-peptide interface highlights the molecular basis for this high-affinity coordination. The N-terminal segment, containing Gln46 and Arg47 previously identified as key for binding in biochemical assays (Fig. 1b), occupy the entrance of the HSL3 binding pocket and anchor the peptide through a network of direct and water-mediated hydrogen bonds with the receptor (Fig. 2d). Mutation of the receptor residue Gln120, which coordinates this N-terminal peptide segment, to alanine reduced binding affinity by ∼ 60-fold, highlighting its functional relevance in peptide coordination. In the core region, additional polar interactions including those with Ser53, a residue conserved in HAE-IDL complexes, Asn56 and Arg51 further stabilise the receptor-peptide interface (Fig. 2d,f). At the C-terminal end, the disulfide-stabilised cyclic loop fits into a previously uncharacterised cavity of HSL3, where an extended network of hydrogen bonds and hydrophobic contacts, reinforces peptide binding and confers overall structural stability (Fig. 2d). Within this region, CTNIP4 Ile61 plays a critical role by occupying a defined hydrophobic pocket in the receptor formed by Gly382 and surrounding residues. Substitution of Ile61 to tyrosine severely compromises this interaction and reduces binding affinity by ∼120-fold (Fig. 2 d,e). In contrast, mutations of Asn60, which forms an extended hydrogen-bond network with HSL3 residues Tyr386, Gln408, and Glu360, has no dramatic effect on peptide binding to the receptor. Additional receptor residues in the core and C-terminal pocket, notably Asp288 and Arg358, contribute to the stabilisation of the cyclic loop, further locking the peptide into place within the binding groove (Fig. 2d,f). Notably, the overall topology of the HSL3 ligand-binding pocket diverges from that of the HAE-HSL receptors and other closely related LRR membrane proteins, revealing a previously uncharacterised peptide-binding mode among plant RKs (Supplementary Fig. 4). In contrast to HAE-HSL receptors, whose pocket tapers toward the C-terminal region to accommodate the linear IDA/IDL peptides^6,7^, HSL3 displays an expanded C-terminal pocket that forms a distinct hydrophobic cavity shaped to accommodate the disulfide-locked cyclic architecture of CTNIP4. This structural extension generates a unique recognition surface that enables HSL3 to stabilise and coordinate the cyclic CTNIP4 fold, a feature absent from its sequence-related paralogues (Supplementary Fig. 4). From the ligand perspective, CTNIPs and IDA/IDL peptides share a broadly similar N-terminal and core region but diverge markedly at their C-terminal end. The additional residues and disulfide-stabilised loop in CTNIPs introduce a structural determinant that promotes selective docking into the HSL3 pocket, thereby preventing cross-recognition by other IDA-type receptors and *vice versa* (Suplementary Fig. 5)^4^. Together, these structural insights define HSL3 as a specialised receptor for cyclic CTNIP peptides, illustrating how subtle architectural variations within conserved LRR-RK frameworks can drive ligand specificity and define a new paradigm for peptide perception in plants.

**Fig 2.**
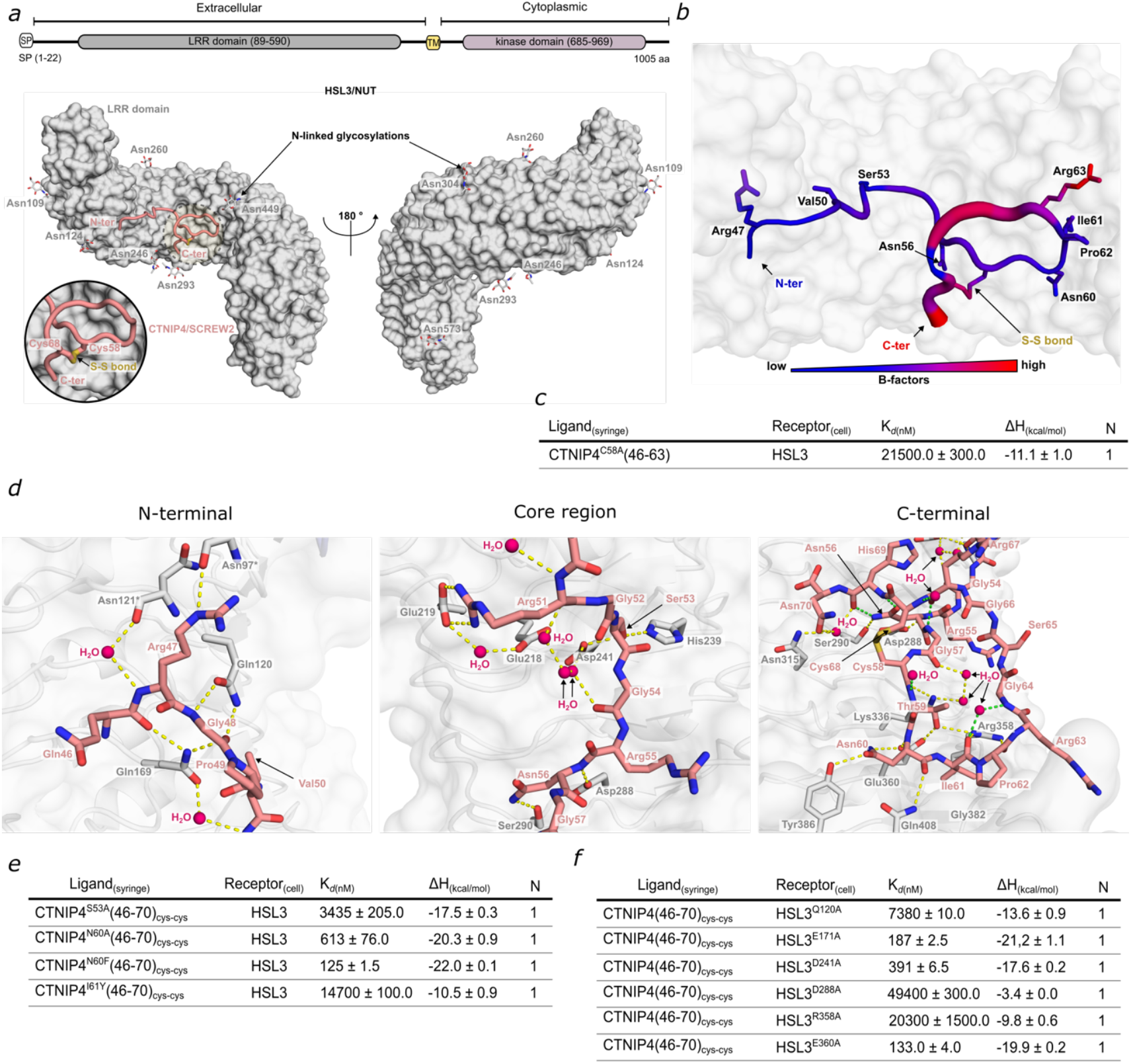
HSL3 pocket recognizes the unique disulfide-stabilised fold of CTNIP4. **a**, Domain architecture of HSL3, comprising a signal peptide (SP), an LRR ectodomain, a single transmembrane helix (TM), and a cytoplasmic kinase domain. Surface representation of HSL3 ectodomain (grey) bound to CTNIP4 (pink cartoon) shows the peptide adopting a cyclic conformation stabilised by an intramolecular disulfide bond. The disulfide bond is shown in yellow, and N-linked glycans are displayed as grey sticks. **b**, Temperature factor (B-factor) representation of CTNIP4 within the HSL3 binding pocket, illustrating receptor-peptide coordination. Regions of low B-factors correspond to tightly-coordinated peptide segments stabilised by receptor contacts. **c**, The disulfide-locked cyclic C-terminal region of CTNIP4 is essential for high-affinity HSL3 binding. Isothermal titration calorimetry (ITC) of the truncated CTNIP4(46-63) linear peptide vs HSL3. **d**, Detailed view of CTNIP4 coordination along the HSL3 binding pocket. HSL3 is shown in grey, CTNIP4 in pink sticks, and water molecules in hot pink spheres. Hydrogen bonds between receptor and peptide are depicted as yellow dashed lines, and intramolecular polar contacts stabilising the cyclic peptide fold are shown in green dashed lines. **e**,**f**, ITC analyses of engineered CTNIP4 and HSL3 variants reveal key residues contributing to receptor-peptide coordination. Tables summarise dissociation constants (Kd, nM), stoichiometry (N = 1), and mean ± SD from at least two independent experiments.

To investigate how CTNIP4 binding promotes receptor activation, we compared the crystal structure of the HSL3-CTNIP4 complex with AlphaFold3-predicted models of the ternary HSL3-CTNIP4-SERK assembly. Structural alignment revealed a similar receptor-peptide interface geometry (RMSD = 0.53 Å), indicating that CTNIP4 adopts a similar conformation in the binary and ternary complexes (Fig. 3a and Supplementary Fig. 6). In the modelled ternary complex, the C-terminal cyclic loop of CTNIP4 interacts with the N-capping domain of the SERK co-receptor, acting as a molecular glue. The ternary assembly is further stabilised through a zipper-like interface formed between HSL3 LRRs 15-21 and the SERK LRRs 1-5, extending to the SERK C-terminal cap domain and establishing a continuous receptor-co-receptor surface (Fig. 3a and Supplementary Fig. 7). Structural comparison of the modelled HSL3-CTNIP4-SERK ternary complex with the experimentally determined HSL1-IDL2-SERK1 structure revealed a similar co-receptor positioning (RMSD = 1.14 Å) and overall conserved complex architecture, but a distinct peptide-mediated activation mechanism in HSL3. Upon CTNIP4 binding, the receptor-peptide interface reorganises to generate a new predominantly hydrophobic surface along the C-terminal region of the peptide guided by the disulfide bond. This newly formed apolar patch directly engages the N-terminal cap of the SERK co-receptor, acting as a molecular bridge that stabilises the ternary assembly (Fig. 3c). Isothermal titration calorimetry confirmed that while HSL3-CTNIP4 binding is enthalpy-driven, subsequent co-receptor recruitment is predominantly entropy-driven, consistent with the burial of this hydrophobic interface (Supplementary Fig. 7). Mutations that disrupt the disulfide-stabilised cyclic fold or the local peptide region recognised by SERKs-(CTNIP4^I61Y^ and CTNIP4^N60A/F^) markedly reduced co-receptor recruitment and overall binding affinity relative to the wild-type CTNIP4(46-70)_Cys-Cys_ peptide (Fig. 3d). Likewise, truncated or linear variants lacking the cyclic loop (CTNIP4^C58A^(46-63) and CTNIP4^C58A.C68A^(46-70)) exhibited 20-60-fold weaker binding, highlighting the importance of the cyclic conformation and its hydrophobic environment for co-receptor recognition and receptor activation (Fig. 2d and 3d). Consistent with this model, substitution of key HSL3 residues forming the receptor-co-receptor interface (Arg452Tyr and Leu547Tyr) completely abolished SERK1 association *in vitro*, validating the structural model and confirming the critical role of this zipper-like hydrophobic interface in stabilising the active ternary complex (Fig. 3a,e and Supplementary Fig. 1 and Fig. 7).

**Fig 3.**
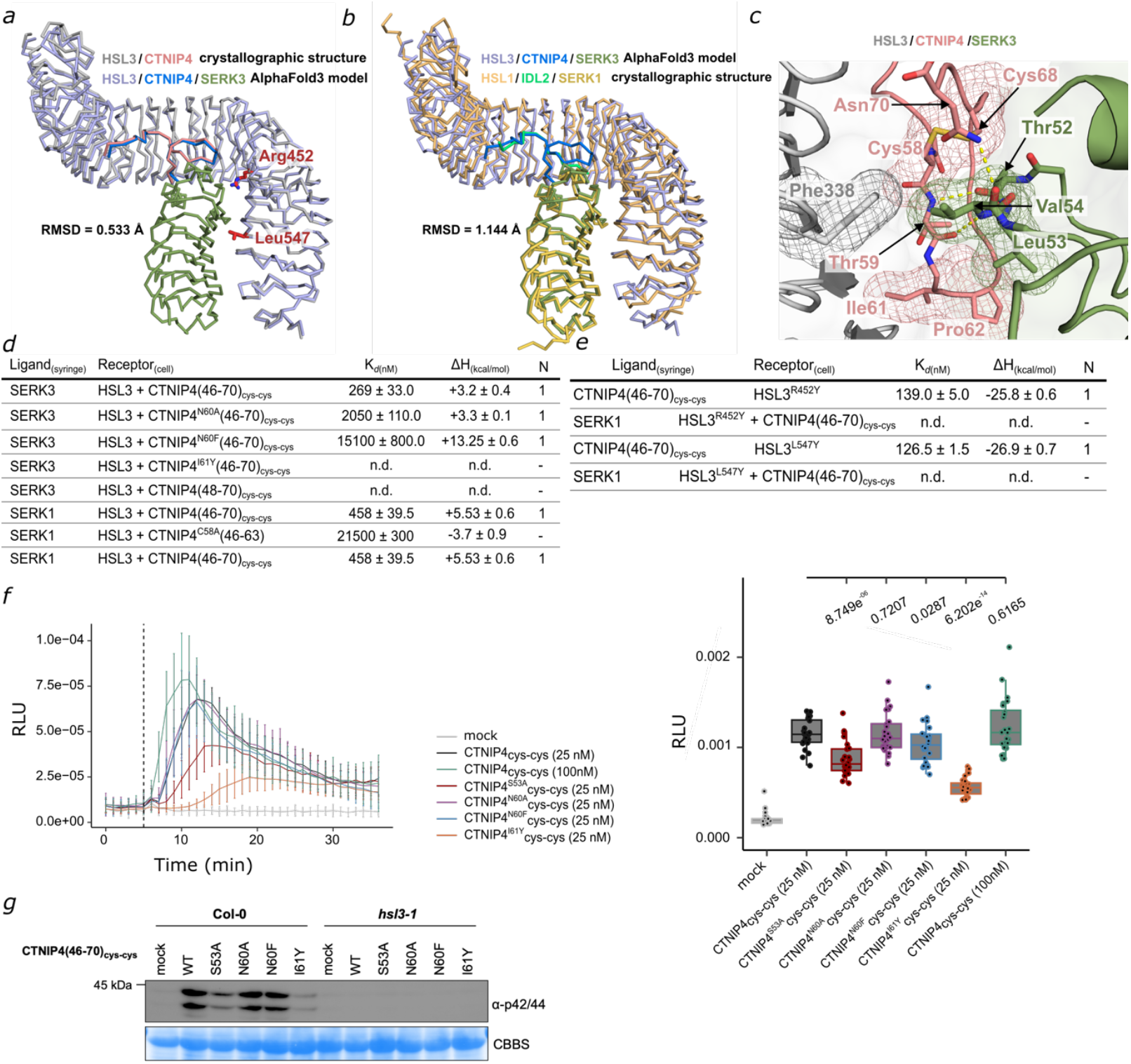
HSL3-CTNIP4 complex defines a new phytocytokine recognition and receptor activation interface. **a**, Structural superimposition of the HSL3-CTNIP4 complex (PDB ID:9T26) with the AlphaFold model of the ternary HSL3-CTNIP4-SERK3 complex. Residues Arg452 and Leu547 at the receptor-co-receptor zipper-like interface are highlighted as red sticks. RMSD=0.533 Å. **b**, Structural comparison of the AlphaFold-predicted HSL3-CTNIP4-SERK3 complex with the experimentally determined HSL1-IDL2-SERK1 complex (PDB ID: 7OGQ). RMSD=1.144 Å. Structures are shown in ribbon representation. **c**, The disulfide-locked cyclic conformation of CTNIP4 bound to HSL3 generates a hydrophobic surface recognised by the N-terminal region of SERK co-receptors, with SERK3 Val54 embedded into the receptor-peptide hydrophobic pocket. Close-up view showing the C-terminal cyclic fold of CTNIP4 (pink sticks) interacting with HSL3 (grey) and the SERK co-receptor (green). Polar contacts further stabilising the ternary complex are depicted in yellow dashes. Hydrophobic residues are highlighted by a mesh. **d**, Isothermal titration calorimetry (ITC) analyses of HSL3 binding to CTNIP4 variant peptides that alter the interface with the SERK co-receptors. **e**, ITC assays of HSL3 mutants targeting the receptor-co-receptor zipper-like interface using the folded CTNIP4(46–70) peptide. ITC tables summarise dissociation constants (Kd, nM), stoichiometry (N = 1), and mean ± SD from at least two independent experiments; n.d., no binding detected. **f**, Calcium influx responses in relative luminescence units (RLUs) triggered by CTNIP4(46-70)_Cys-Cys_ interface mutants in wild-type Col-0^AEQ^ seedlings. Peptide concentration used was 25 nM except if indicated differently. Data represent mean ± SD (n = 12), the vertical dotted line indicates the timing of the treatment. Considering the cumulative RLUs (3-30 min), the box plots indicate the median and the interquartile range (IQR), the whiskers extend to the most extreme data points within 1.5 times the IQR. Significance was tested by performing non-parametric Wilcoxon-Mann-Whitney tests between CTNIP4(46-70)_cys-cys_ and CTNIP4(46-70)_cys-cys_ variants. **g**, MAP kinase activation in response to 1 nM folded CTNIP4(46-70)_Cys-Cys_ interface variant peptides. Treatment was given for 20 mins. Data shown are from one representative experiment out of three with similar results.

To translate these structural and thermodynamic findings into a physiological context, we assessed signaling competence of the receptor and peptide variants *in planta* using cytosolic Ca^2+^ influx and MAP kinase activation assays. CTNIP4 mutants that impaired either peptide-receptor or receptor-co-receptor interfaces exhibited a reduced Ca^2+^ influx or MAPK phosphorylation (Fig. 3f,g), mirroring their diminished binding affinities *in vitro*. These results establish that the disulfide-stabilised cyclic architecture of CTNIP4 and the corresponding hydrophobic receptor-co-receptor interface are essential for productive complex formation and downstream signaling activation.

## Discussion

Precise peptide recognition by structurally similar LRR-RKs is essential for coordinating diverse plant responses. Our study reveals that HSL3/NUT selectively recognizes CTNIP/SCREW peptides through their N-terminal anchoring region and the disulfide-locked C-terminal cyclic loop stabilized by the Cys58–Cys68 bond. HSL3’s C-terminal ligand-binding pocket exhibits a unique hydrophobic cavity accommodating the cyclic loop, distinguishing HSL3 from IDA/IDL-binding and other peptide LRR-RKs^6,7^. SERK co-receptor recruitment occurs mainly via a unique hydrophobic interface generated upon peptide binding, leading to an entropy-driven receptor activation. Disruption of the cyclic fold or hydrophobic interface abolishes ligand-induced signaling responses. These findings establish a new paradigm for peptide-mediated receptor activation, demonstrating how subtle structural adaptations within conserved LRR scaffolds confer ligand specificity and orchestrate receptor-co-receptor assembly in plants.

## Data availability

All data are available in the main text or the supplementary materials. The HSL3-CTNIP4 structure is deposited at the Protein Data Bank (PDB ID 9T26). Materials and raw data files are available upon request to the corresponding authors Cyril Zipfel or Julia Santiago.

## Acknowledgements

The authors thank the ESRF for providing beam time and the staff of beamline in Grenoble, for their technical assistance during data collection. This research was supported by the University of Lausanne (JS) and the Swiss National Science Foundation grant no. 310030_204526 (JS). the University of Zurich (CZ), the Gatsby Charitable Foundation (CZ), the European Research Council under the European Union’s Horizon 2020 research and innovation programme no. 773153 (project ‘IMMUNO-PEPTALK’) (CZ), and the Swiss National Science Foundation grant no. 10001549 to CZ and JS. Postdoc EMBO fellowship (ALTF 1061-2023) to MO. University of Zurich Postdoc Grant, Uniscientia Stiftung, to SS.

## Author information

These authors supervised this work: Cyril Zipfel and Julia Santiago

## Contributions

JS, PJS, and CZ conceptualized the study. PJS led the experimental work, including protein expression, purification, crystallization, X-ray diffraction, structural analyses, mutant design, and binding assays. OJ performed AF3 modelling and preliminary phenotypic characterization of peptide and receptor variants. SS designed and performed additional Ca^+2^ assays. HCY designed and performed additional MAPK phosphorylation assays. MO designed and performed confocal imaging of HSL3 variants. CB purified HSL3 protein variants and performed SEC and SDS protein analysis. JR analysed data and provided conceptual advice to the project. KWB cloned HSL3 variants and provided conceptual advice to the project. SS designed and performed Ca^+2^ assays. HCYG performed MAPK assays. JS and CZ secured funding, oversaw project administration, and provided supervision throughout the study. The original draft of the manuscript was written by JS, with writing, review and editing contributions from all other authors.

## Methods Synthetic peptides

All peptides used in this study were synthesized and purchased from GenScript. Peptide sequences, including point mutations and the single disulfide bridge, are listed below. The linear production of CTNIP4(48-70)_linear_ was done according to the manufacturer and diluted right before the ITC assays to avoid oxidation.

### Peptide sequences used in this study

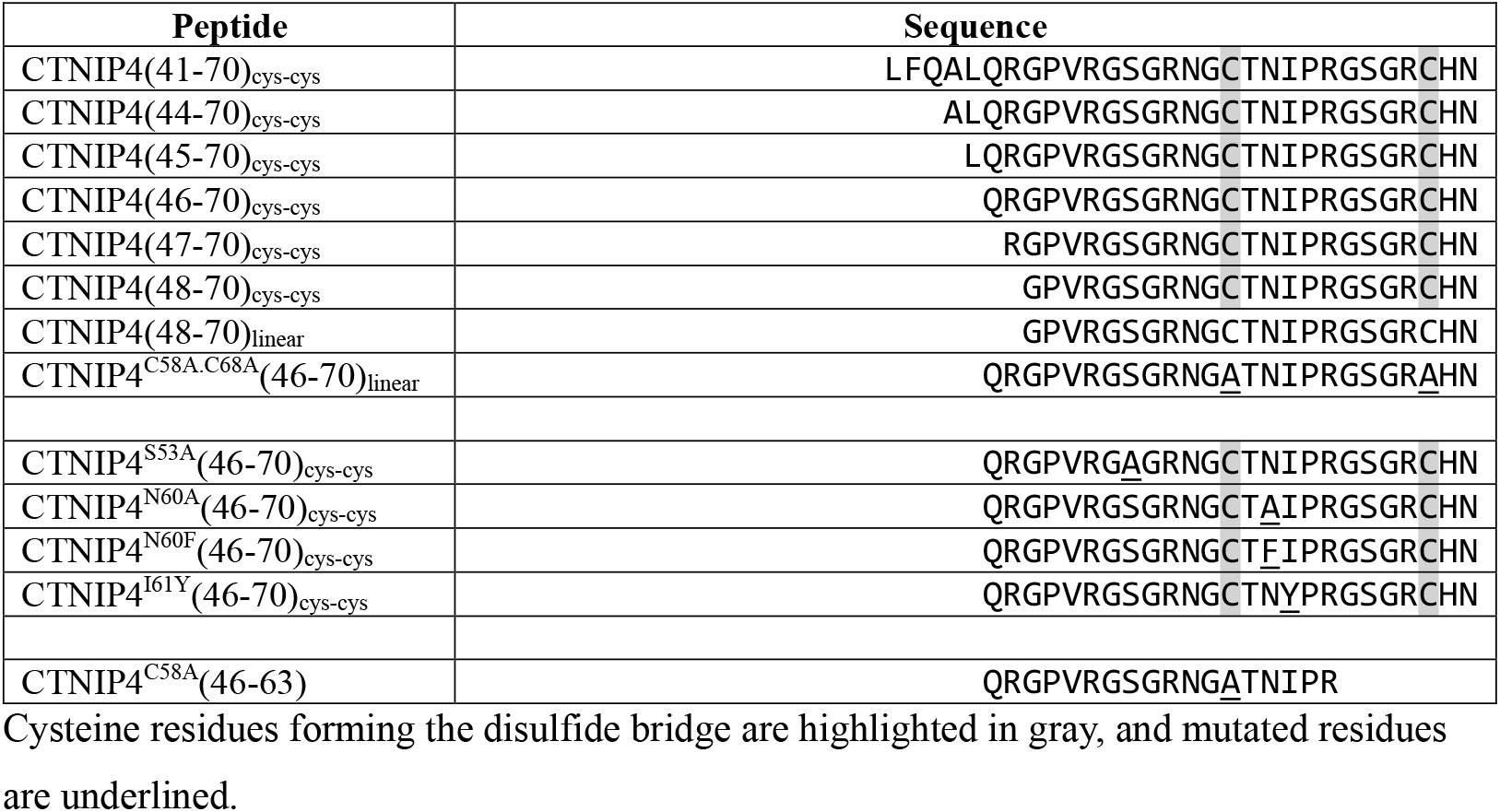

### Protein expression and purification

The wildtype ectodomain of HSL3 (AT5G25930, residues 23–627) and mutants generated through site directed mutagenesis and confirmed by sequencing using the primers indicated in Supplementary Table 2, were cloned into a modified pFastBac donor vector (Geneva Biotech) harbouring the 30K *Bombyx mori* secretion signal peptide^21^, and with a TEV (tobacco etch virus protease) cleavable C-terminal StrepII-9xHis tag. SERK1 (AT1G71830, residues 24– 213) and BAK1 (AT4G33430, residues 1–220) were cloned in the same vector harbouring azurocidin and no signal peptide respectively. Baculovirus vectors were generated in DH10 MultiBac *E. coli* cells (Geneva Biotech). Virus amplification was carried out in *Spodoptera frugiperda* Sf9 cells (Geneart, Thermo Fisher Scientific) and was used to infect *Trichoplusia ni* Tnao38 cells^27^ for protein expression. The cells were grown for 1 day at 28°C and for 2 days at 21°C with gentle shaking. The secreted proteins were subjected to tandem affinity purification, using Ni^2+^ (HisTrap excel, equilibrated in 25 mM KPi, and 500 mM NaCl, pH 7.8; Cytiva) and Strep columns (Strep-Tactin Superflow high-capacity; IBA) equilibrated in 25 mM Tris, 250 mM NaCl, 1 mM EDTA, pH 8.0. Affinity tags were removed using His-tagged TEV protease in a 1:50 ratio at 4°C overnight. Separation of cleavage tags and aggregated proteins was performed using size-exclusion chromatography on a Superdex 200 Increase 10/300 GL column (Cytiva) equilibrated in 20 mM sodium citrate, 150 mM NaCl, pH 5.0. Proteins were analysed for purity and structural integrity by SDS-PAGE.

### Crystallization and data collection

Purified HSL3 ectodomain was partially deglycosylated prior to crystallization. HSL3 was incubated with a mixture of Endo H and Endo F1 glycosidases for 3 hours at room temperature at a ratio of 20:1:1 (protein:Endo H:Endo F1) in a buffer containing 100 mM Tris–HCl and 150 mM NaCl, pH 7.5. The partially deglycosylated HSL3 was further purified by size-exclusion chromatography on a Superdex 200 Increase 10/300 GL column (Cytiva) equilibrated in 20 mM sodium citrate, 150 mM NaCl, pH 5.0, and used for crystallization screens.

Crystals of HSL3 in complex with CTNIP4(46-70)_Cys–Cys_ were obtained using the sitting-drop vapor diffusion technique at 18 °C. Crystals were grown in a solution containing 0.1 M sodium acetate, pH 5.5, 0.2 M sodium chloride, and 25% (w/v) PEG 3350, using a 1:1 ratio of protein complex to crystallization buffer (200 nl of each). Diffracting crystals were cryoprotected with mother liquor supplemented with 20% ethylene glycol and snap-frozen in liquid nitrogen. A native, high-resolution X-ray diffraction dataset was collected at 100 K (–173 °C) at the European Synchrotron Radiation Facility (ESRF), Grenoble, France.

### Structure determination and refinement

The structure of HSL3 in complex with the CTNIP4(46-70)_cys-cys_ peptide was solved by molecular replacement (MR) using the AlphaFold^28^ model of HSL3 (AlphaFold ID: AF-Q9XGZ2-F1) as a search model, considering only the ectodomain. Solvent analysis suggested the presence of one HSL3/CTNIP4 complex per asymmetric unit. MR was performed with Phaser within the CCP4 software suite^29^ and manually rebuilt in COOT^30^. The model was refined to 2.12 Å resolution using Refmac5^31^, including automatic TLS refinement, which was applied to achieve optimal R-factors. Water molecules were added where supported by electron density maps and appropriate hydrogen-bonding distances. Model validation was carried out using Coot validation tools and MolProbity^32^, and the refined coordinates have been deposited in the Protein Data Bank under PDB ID: 9T26.

### Isothermal titration calorimetry (ITC)

Experiments were performed at 25 °C using a MicroCal PEAQ-ITC (Malvern Instruments) with a 200 µL standard cell and a 40 µL titration syringe. HSL3 and its variants, together with SERKs (SERK1 and SERK3), were gel-filtrated into the ITC buffer (20 mM sodium citrate, 150 mM NaCl, pH 5.0). Molar protein concentrations were calculated using individual molecular weights and molar extinction coefficients. For reference, HSL3: 68.43 kDa, 56,880 M^−1^ cm^−1^; SERK1: 21.94 kDa, 17,210 M^−1^ cm^−1^; SERK3/BAK1: 21.97 kDa, 17,085 M^−1^ cm^−1^. Values for HSL3 variants were calculated using ProtParam.

A typical experiment consisted of injecting 2 µL of a 140–800 µM solution of CTNIP4 peptides (peptides used are listed above) into a 15 µM solution of HSL3 (or variants) in the cell at 150 s intervals with 500 rpm stirring speed. For SERKs versus HSL3/CTNIP4 peptide experiments, individual SERKs (SERK1 or SERK3) were titrated from the syringe into the cell containing 15 µM HSL3 (or variants) pre-incubated with saturating or near-saturating concentrations of CTNIP4 peptides, using the same injection pattern. ITC data were baseline-corrected using the software’s offset correction.

Experiments were performed in duplicates, and data were analyzed with the same software. All ITC runs used for data analysis had N values ranging from 0.9 to 1.1, which were fitted to 1 during analysis.

### Plant materials and conditions

*Arabidopsis thaliana* ecotype Columbia (Col-0) aequorin (Col-0^AEQ^)^33^ was used for cytoplasmic calcium measurements and were grown in growth chambers (20 °C, 60 % RH and 10:14 light:dark cycles).

For MAPK assay, 10-day-old *Arabidopsis thaliana* Col-0 and *hsl3-1* seedlings were used. Seeds were surface sterilized using chlorine gas for 5-6 h and plated on half-strength Murashige and Skoog (½ MS) media supplemented with 1 % sucrose and 0.9 % agar and were stratified for 2 days in the dark at 4 °C. Seeds were then moved to a growth chamber with conditions 16 h day/8 h night at 22 °C/18 °C respectively. Germinated seedlings were transferred to 24 well plates with two seedlings per well containing 1 mL of liquid ½ MS and grown for 7 days in the growth chamber.

### Cytoplastic calcium measurements in Arabidopsis

Leaf punches were taken with a 4-mm biopsy punch of Arabidopsis aequorin and floated in 100 μL of H_2_O with 20 μM coelenterazine (Merck), using individual cells of a white 96-well bottom plate (Greiner F-Boden, lumitrac, med. Binding, [REF 655075]). After overnight incubation, the coelenterazine solution was replaced with 100 μL H_2_O and rested for circa 30 min in the dark. Luminescence was quantified in a Berthold Tristar 3 plate reader every minute for 5 min before and 30 min post elicitor treatment using an integration time of 250 ms. R and the R-packages dplyr (v1.1.2), ggpubr (v.0.6.0), and ggplot2 (v3.4.2) were used to analyze and plot the data. The resulting figure was edited in Corel-DRAW Home & Student x7.

### MAPK activation assays

For MAPK assay, plant samples were ground into fine powder in liquid nitrogen homogenized in protein extraction buffer (50 mM Tris-HCl pH 7.5, 100 mM NaCl, 10% glycerol, 2 mM EDTA, 5 mM dithiothreitol, 1% IGEPAL CA630, 2 mM sodium molybdate, 1 mM sodium fluoride, 1 mM sodium orthovanadate, 4 mM sodium tartrate, protease inhibitor cocktail). Samples were incubated with rotation at 4 °C for 30 min and then centrifuged at 16,000 *g* for 15 min. Supernatants were collected and SDS loading buffer was added for western blotting. Proteins were resolved using 10 % SDS-PAGE and transferred to PVDF membrane. Membranes were blocked using 5 % non-fat milk and probed for MAPK activation using p44/42 MAPK (Cell Signaling, 9102, RRID: AB_330744, 1:1,000) primary antibody and HRP-conjugated anti-rabbit IgG-peroxidase (Sigma, A0545, RRID: AB_257896, 1:10,000) secondary antibody. Protein bands were detected using enhanced chemiluminescence (ECL) substrate (Thermo Scientific, 32106 or 34096). Coomassie brilliant blue solution was used to detect protein loading control.

