## Supplementary Information for "Disulfide bond sculpts a peptide fold that mediates phytocytokine recognition"

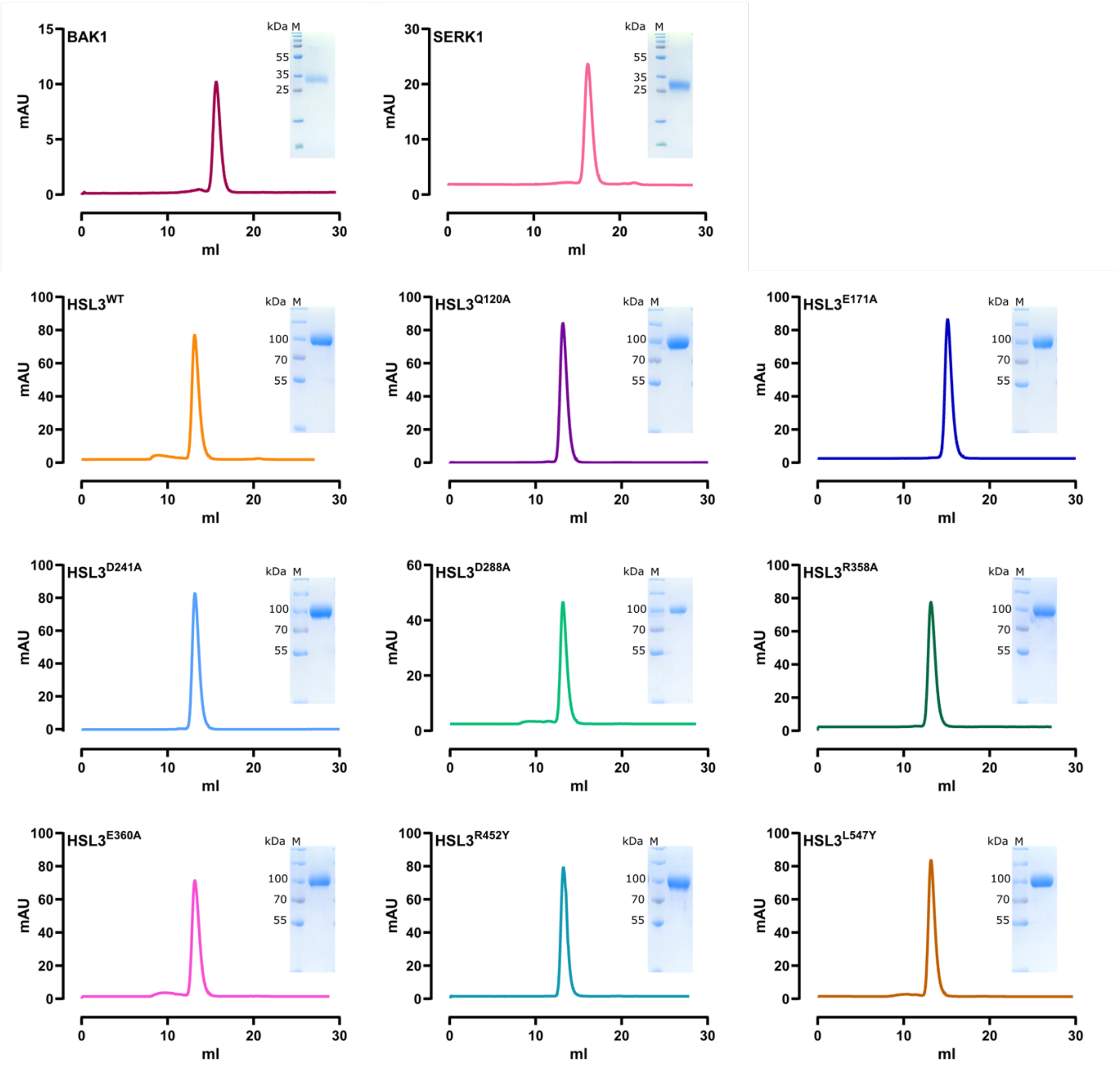

**Supplementary Fig. 1. Purification profiles of HSL3 wild-type, HSL3 variants, and SERKs co-receptor proteins**. Size-exclusion chromatography (SEC) traces and SDS–PAGE analyses of insect cell–expressed and purified HSL3 wild-type, HSL3 variant, and SERK co-receptor ectodomains.

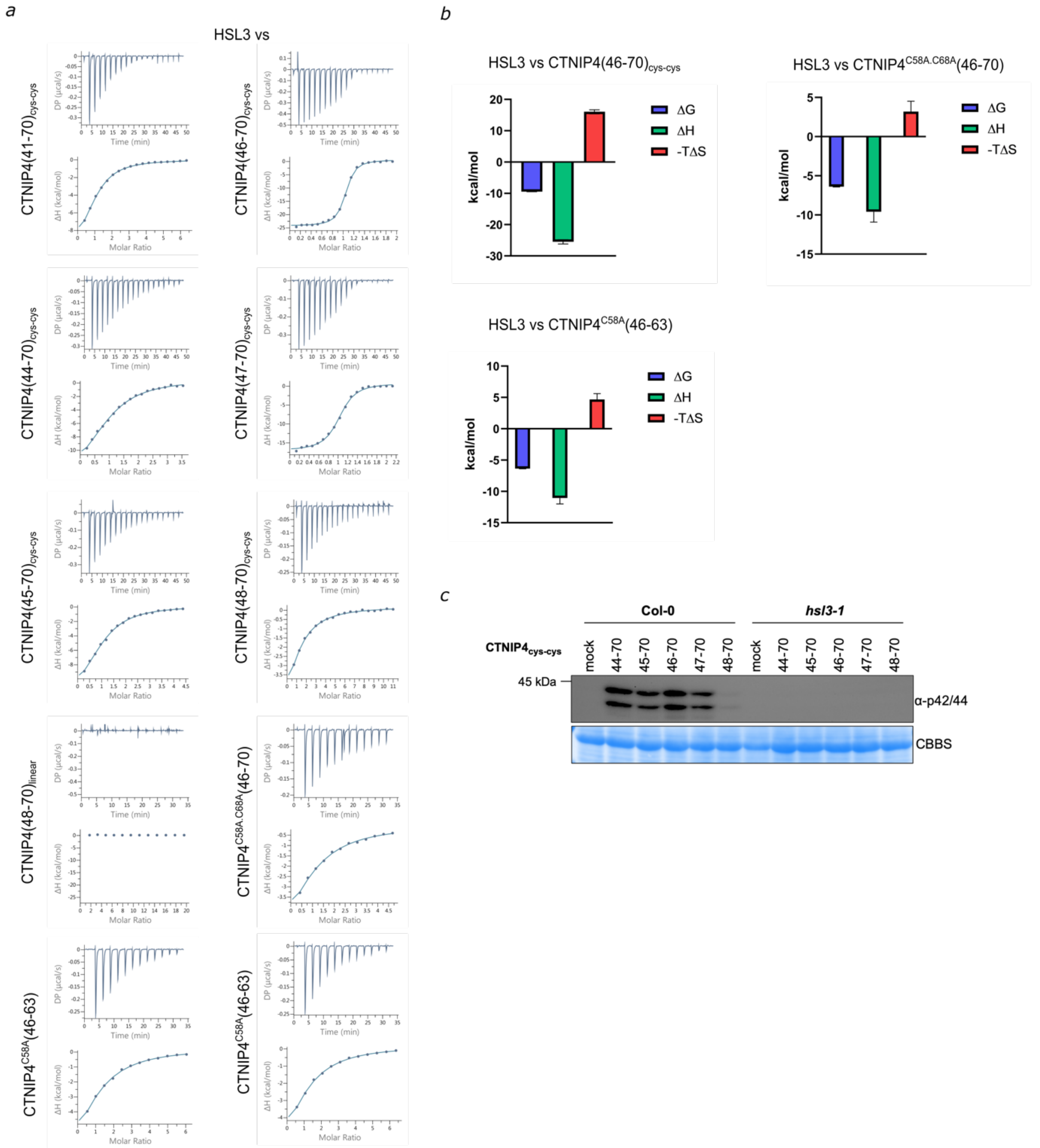

**Supplementary Fig. 2. ITC experiments and thermodynamic parameters of CTNIP4 variants and MAPK activation replicate of experiments shown in Fig. 1. a,** ITC thermograms of independent replicate experiments corresponding to those shown in Fig. 1b. **b**, Thermodynamic profiles of CTNIP4 peptides binding to HSL3 derived from ITC measurements, showing changes in Gibbs free energy (ΔG), enthalpy (ΔH), and the entropic contribution (−TΔS). **c**, Independent replicate of MAP kinase (MAPK) activation assays shown in Fig. 1d.

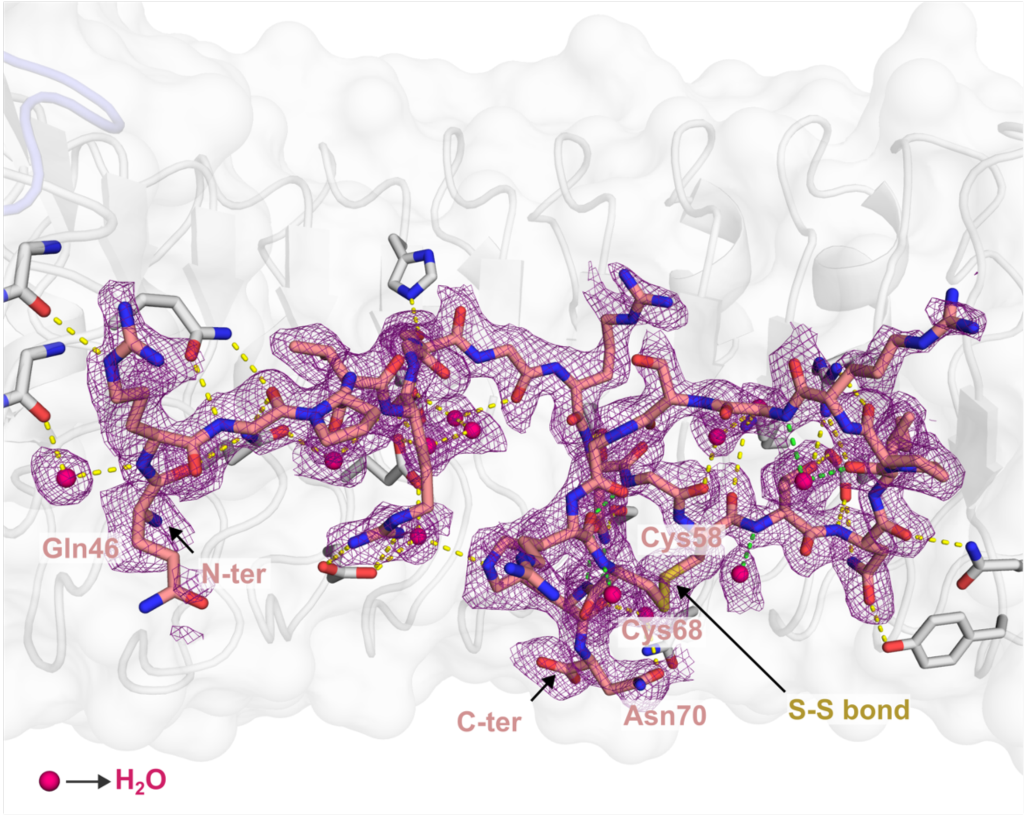

**Supplementary Fig. 3. CTNIP4 omit map within HSL3 binding pocket.** Electron-density omit map (|Fo|–|Fc|), contoured at 1.5σ (purple mesh), highlights the CTNIP4 peptide (pink sticks) and crystallographic water molecules (hot pink spheres) participating in the direct polar interaction network (yellow dashed lines). Peptide intramolecular interactions are depicted in green dashes. The HSL3 receptor is shown in grey. Disulfide-linked residues Cys58 and Cys68 constrain CTNIP4 into a cyclic conformation accommodated within the C-terminal ligand-binding pocket of HSL3.

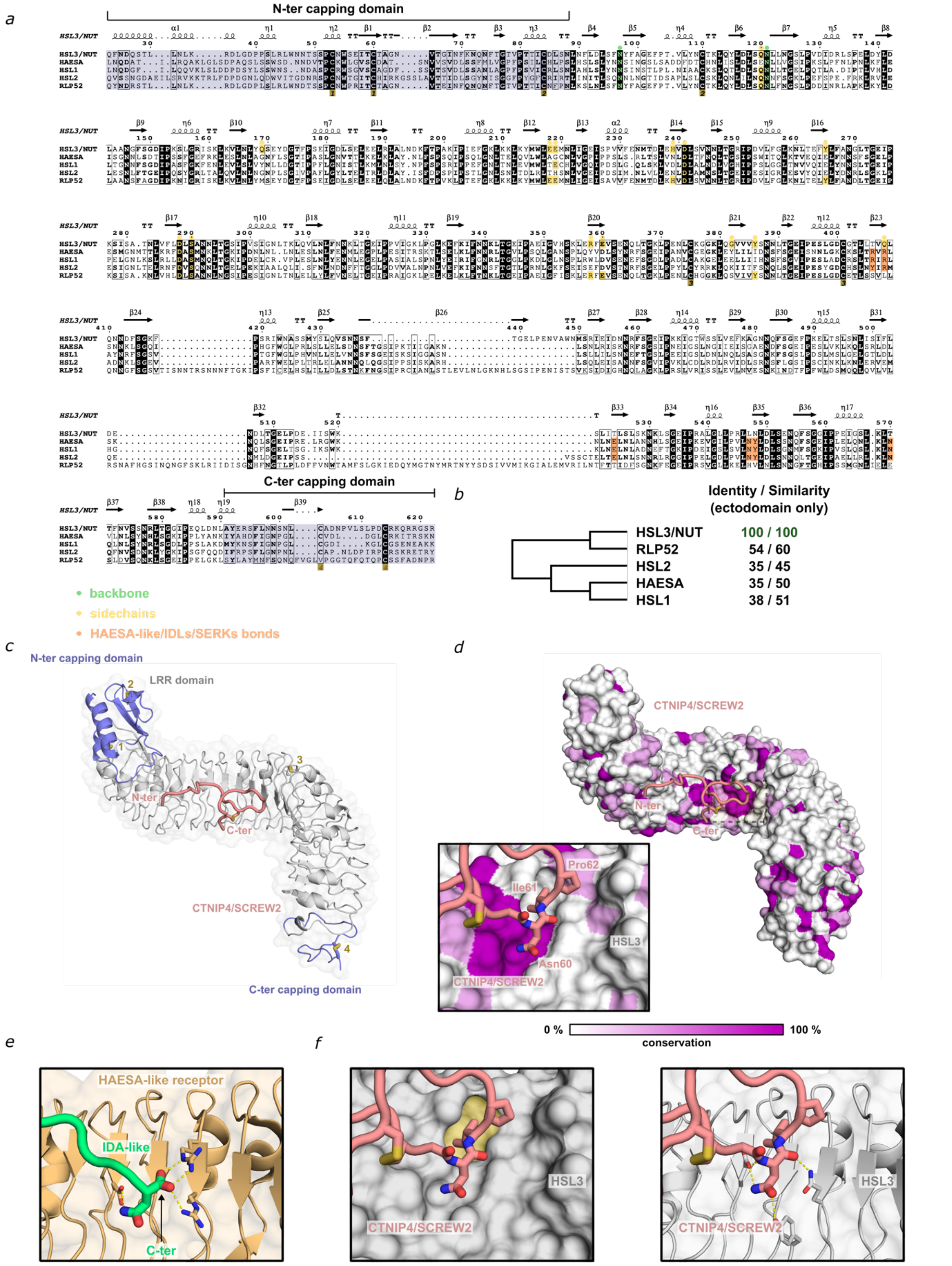

**Supplementary Fig. 4. HSL3 has evolved a distinct C-terminal pocket architecture to selectively engage the cyclic, disulfide-stabilised CTNIP peptides. a,** Comparison of HSL3 with related LRR-RKs reveals diversification of the C-terminal binding pocket, which underlies the receptor's specificity for the cyclic, disulfide-stabilised CTNIP peptide fold. Ectodomain sequence alignment of HSL3 with its closest LRR relatives is shown, including secondary structure elements and residue numbering based on the HSL3 crystal structure (PDB ID: 9T26, this study). Residues involved in backbone and side-chain interactions with CTNIP4 are highlighted. Links to HAESA-like receptors, IDL peptides, and SERK co-receptors are indicated to illustrate structural differences that define *bona fide* ligand recognition. **b**, Phylogenetic analysis of HSL3 and closely related LRR proteins, with sequence identity scores relative to HSL3 shown. **c**, Topology of the HSL3-CTNIP complex, showing that HSL3 contains the canonical N-terminal and C-terminal capping domains sandwiching a solenoid composed of 21 LRRs. Conserved disulfide bonds are indicated in yellow. The CTNIP4 peptide binds to the inner surface of the LRR receptor. **d**, Surface conservation analysis of HSL3 and related LRRs from panel b, illustrating the differences in pocket architecture. **e**, Close-up view of the C-terminal ligand pocket in HAESA-like receptors coordinating the linear IDA peptide via a conserved pair of Arg residues that constrict the binding groove. **f**, Detailed view of the HSL3 C-terminal pocket in cartoon and surface representation, showing how it accommodates the cyclic loop of CTNIP4. The extended C-terminal end of HSL3 forms a unique hydrophobic cavity that, together with polar contacts, anchors the disulfide-stabilised peptide fold and enables selective peptide recognition.

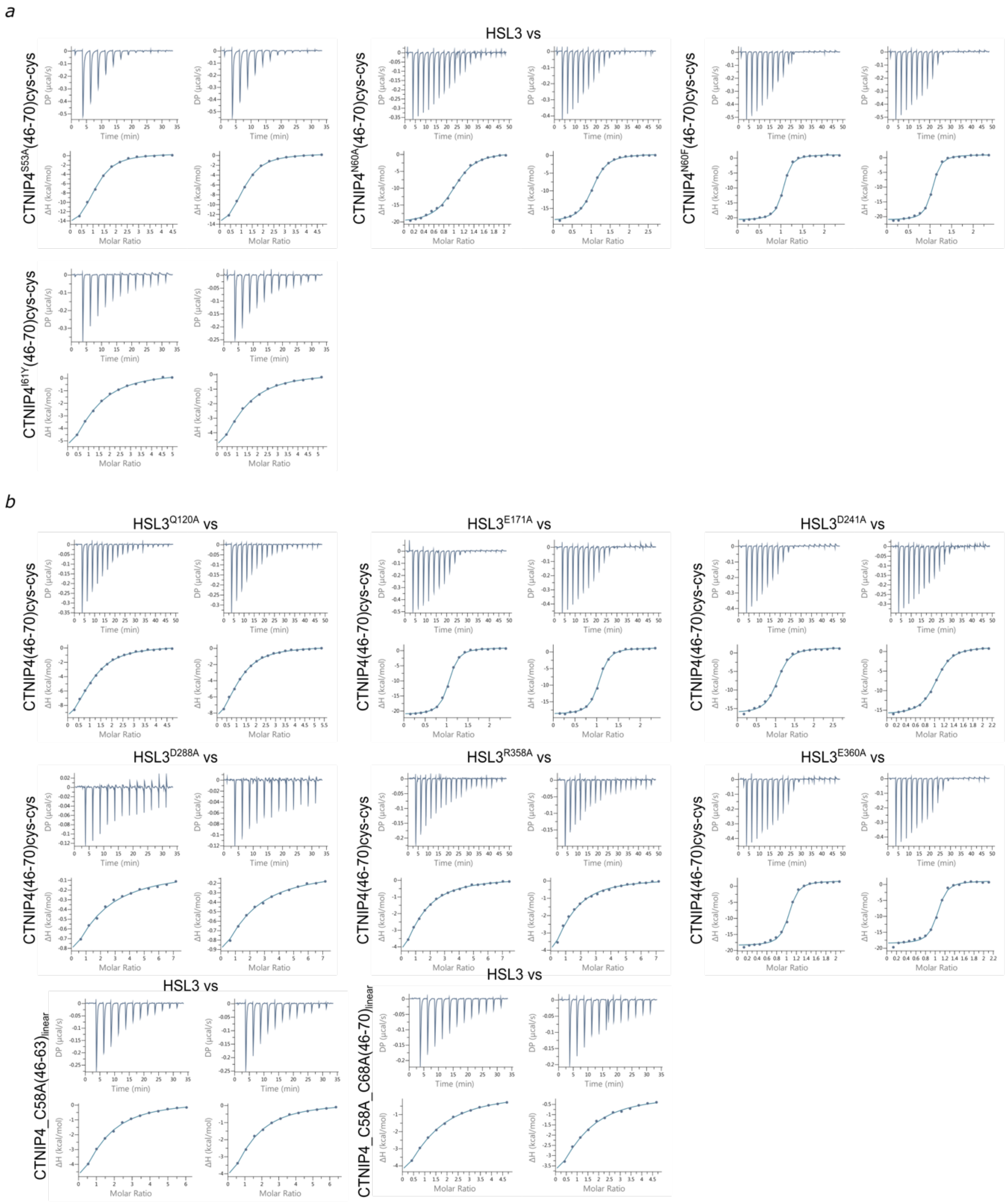

**Supplementary Fig. 5. ITC experiments of HSL3 wild-type and variants vs CTNIP4 peptide.** ITC thermograms of independent replicate experiments corresponding to those shown in Fig. 2c,f.

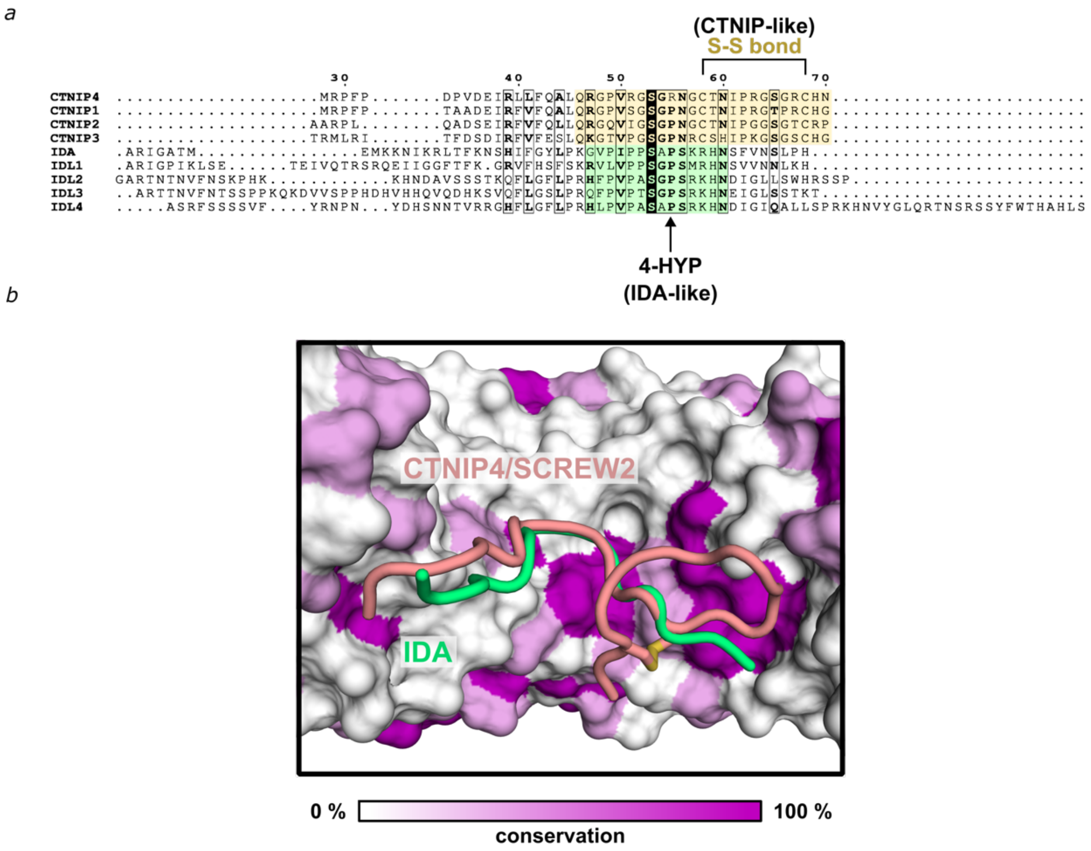

**Supplementary Fig. 6. Distinct peptides have evolved a different receptor recognition mechanism. a,** Sequence alignment of CTNIPs and IDA/IDLs. In green are depicted the mature processed IDA/IDL peptides. The hyproxylated Pro residues in indicated. In yellow are highlighted the minimal higher affinity CTNIPs mapped in this study. Disulfide bond stabilising the C-ter part of the peptide is indicated. **b**, Detail view of the pocket surface conservation of HSL3 and related LRR-RKs. The structure of HSL3-CTNIP4 is superimposed to HSL1-IDL2 (PDB: 7OGQ) to illustrate that the C-ter extension stabilised by the disulfide-bond of CTNIPs, provides a new receptor recognition mechanism (Fig.3b and Supplementary Fig. 4).

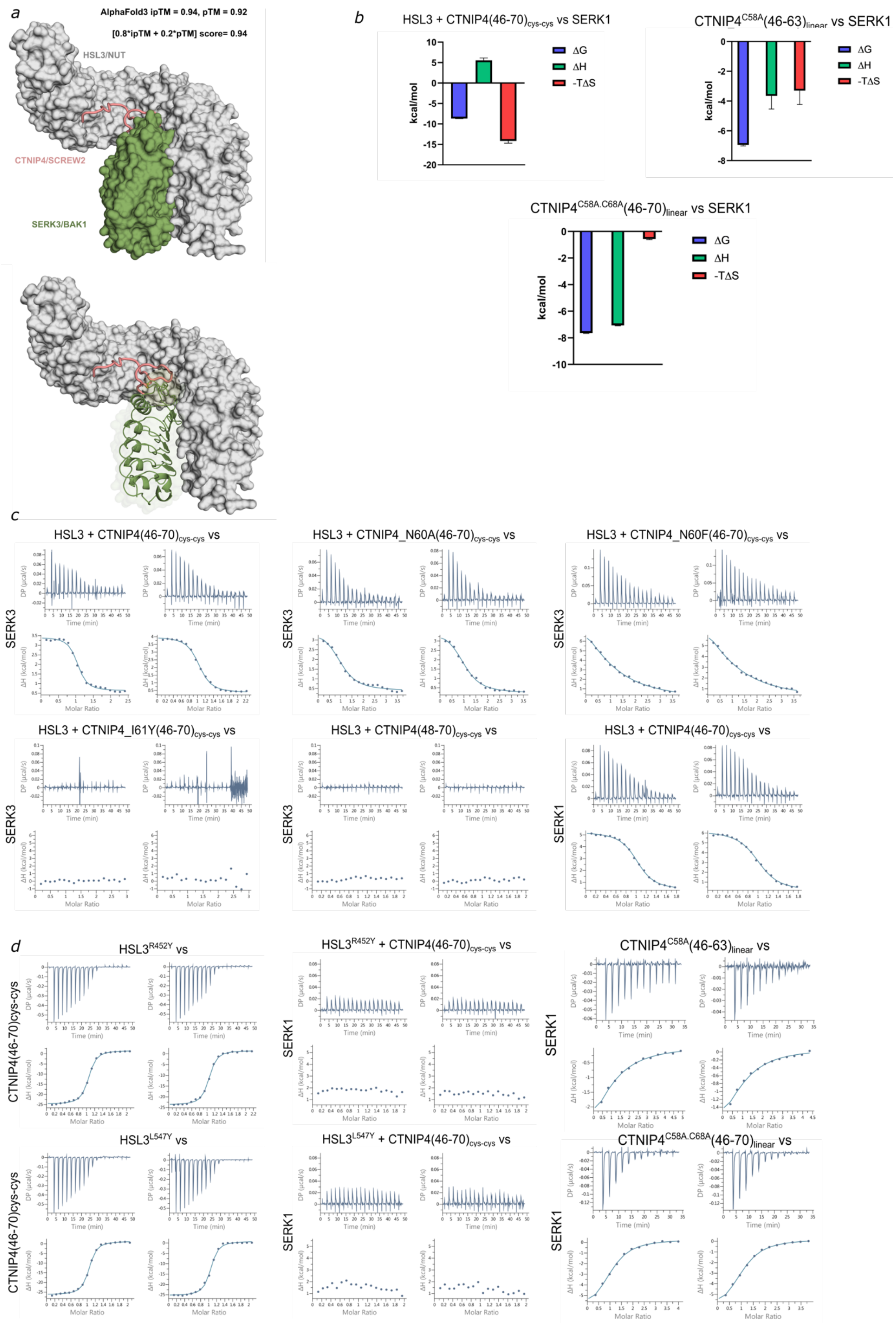

**Supplementary Fig.7. ITC experiments of HSL3-CTNIP4 complex and SERK co-receptors interface shown in Fig. 3. a**, Overall architecture in surface and cartoon representation of the AlphaFold Model HSL3-CTNIP4-SERK3. **b**, Thermodynamic profiles of SERKs binding to HSL3-CTNIP4 derived from ITC measurements, showing changes in Gibbs free energy ($\Delta G$), enthalpy ($\Delta H$), and the entropic contribution ($-T\Delta S$). The interaction is entropy driven. The enthalpy cost is explained by the displacement of water molecules due to the predominantly hydrophobic nature of the interaction between the receptor-peptide interface and the co-receptor (Fig. 2d, 3c and Supplementary Fig. 3). **c**, ITC thermograms of independent replicate experiments corresponding to those shown in Fig. 3b

### Supplementary Table 1. Crystallographic table of the complex HSL3-CTNIP4.

**Data collection and refinement statistics (molecular replacement)**

| | PDB ID: 9T26 |
|---|---|
| **Data collection** | |
| Space group | C 2 2 21 |
| Cell dimensions | |
| $a$, $b$, $c$ (Å) | 56.69, 131.09, 220.99 |
| α, β, γ (°) | 90.00, 90.00, 90.00 |
| Resolution (Å) | 110.74 - 2.12 (2.18 - 2.12)* |
| $R_{sym}$ or $R_{merge}$ | 0.07(0.64) |
| $I / \sigma I$ | 9.1(1.2) |
| Completeness (%) | 98(97.7) |
| Redundancy | 4.1(4.2) |
| **Refinement** | |
| Resolution (Å) | 110.74 - 2.12 |
| No. reflections | 45998 |
| $R_{work} / R_{free}$ | 0.19 / 0.23 |
| No. atoms | 10044 |
| Protein | 9,371 |
| Ligand/ion | 447 |
| Water | 226 |
| $B$-factors | 58.05 |
| Protein | 56.34 |
| Glycosylations | 99,76 |
| Water | 52.32 |
| R.m.s. deviations | |
| Bond lengths (Å) | 0.016 |
| Bond angles (°) | 2.061 |

*Single crystal diffraction. *Values in parentheses are for highest-resolution shell.

**Supplementary Table 2**. Primers used to generate HSL3 variants for recombinant protein expression in insect cells.

| **HSL3 mutant** | **Primers** |
|---|---|
| Gln120Ala | Fw→GACTTGTCCGCGAACCTGCTCAACGGTTCC<br>Rv→GAGCAGGTTCGCGGACAAGTCCAGGTACTG |
| Glu171Ala | Fw→TACCAGTCCGCGTACGACGGTACTTTCCCA<br>Rv→ACCGTCGTACGCGGACTGGTACAGGTTCAG |
| Asp241Ala | Fw→GAACACGTGGCCCTCTCCGTGAACAACCTG<br>Rv→CACGGAGAGGGCCACGTGTTCCAGGTCGGT |
| Asp288Ala | Fw→GTGTTCCTCGCCTTGTCCGCTAACAACTTG<br>Rv→AGCGGACAAGGCGAGGAACACCAGGTTGGT |
| Arg358Ala | Fw→AAGTTGGAGGCTTTCGAGGTGTCCGAGAAC<br>Rv→CACCTCGAAAGCCTCCAACTTGGAGTGGAC |
| Glu360Ala | Fw→GAGCGTTTCGCGGTGTCCGAGAACCAGCTG<br>Rv→CTCGGACACCGCGAAACGCTCCAACTTGGA |
| Arg452Tyr | Fw→AACATGTCCTATATCGAGATCGACAACAAC<br>Rv→GATCTCGATATAGGACATGTTCCAGGCCAC |
| Leu547Tyr | Fw→CCTCGTCTGTATAACCTGGATCTGAGCGAA<br>Rv→ATCCAGGTTATACAGACGAGGCAGCAAGCC |
